# A developmental transition from early neural hyperexcitability in zebrafish

**DOI:** 10.1101/2025.09.02.673888

**Authors:** Pushpinder Singh, Greeshma Sudha, KS Aiswarya, Anagha Prakash, Mayanglambam Suheshkumar Singh, Amrutha Swaminathan

## Abstract

Early brain development is characterized by plastic phases that play a crucial role in the establishment of functional connectivity. However, this plasticity also renders the developing brain highly sensitive to perturbations. The dynamics of the transition of the immature brain from a hyperexcitable state to a relatively stable network remains poorly understood. Using a combination of brain-wide activity mapping, behavioural and molecular analysis in response to pentylenetetrazole (PTZ), a GABA_A_ receptor antagonist, we assessed neural hyperexcitability at different developmental stages in larval zebrafish and observed a distinct age-dependent trajectory. Specifically, we observed that early stages (4-6 dpf) exhibited high susceptibility to external perturbations, whereas by 9 dpf, both neural activity and behavioral responses were markedly attenuated. Further, upon chronic insult, rather than a cumulative increase in excitability, early life hyperexcitation events rendered the larvae resistant to subsequent induction. Put together, our work maps the temporal transition of the vertebrate brain from a state of hyperexcitability to a phase of relative resistance and has potential implications on understanding the progression of neurological conditions in the pediatric population.

## Introduction

Vertebrate development consists of highly conserved spatiotemporal programs that shape various biological processes [1]. However, rather than a rigidly hardwired program, it comprises phases of growth marked by profound plasticity which render the organism more vulnerable to even transient disruptions, and can have lasting consequences [2,3].

Several developmental processes, including delayed maturation of inhibitory systems, disrupted ion homeostasis, and altered expression of excitatory receptors, render the immature brain more prone to hyperexcitability [4-7]. However, despite the evident importance of early brain development, our mechanistic understanding of the susceptible nature of the immature nervous system remains limited. This is primarily due to the limited access to *in utero* stages in mammalian models, making real-time visualization and experimental manipulation of the brain challenging.

In this study, we performed a detailed temporal mapping of the transitioning of the immature vertebrate brain out of its early hyperexcitable state, when it begins to acquire homeostatic resistance to external insults. We employed the zebrafish (*Danio rerio*), a vertebrate with substantial homology to the mammalian brain, and the advantages of optical transparency, external fertilization and rapid *ex utero* brain maturation, to trace the trajectory of hyperexcitability during brain development [8,9]. Using a combination of brain-wide neural activity, behavioural and molecular readouts, we analyzed the response of larval zebrafish to the GABA_A_ receptor antagonist, pentylenetetrazole (PTZ), during its early development period. Our findings identify a previously unrecognized developmental window during which the immature zebrafish brain transitions from a state of pronounced hyperexcitability towards relative resistance. This shift is evident at both functional and molecular levels and coincides with the reported neural hallmarks such as plateau of the synaptogenesis window [10].

The developing vertebrate brain is uniquely susceptible to a wide range of insults, with disruptions during key developmental windows resulting in conditions such as epilepsy, cerebral palsy, autism spectrum disorders, and cognitive delays [11]. Our identification of a critical developmental phase of transition out of hyperexcitability offers a valuable platform for dissecting the mechanisms that govern homeostatic plasticity and for addressing broader questions about how this dynamicity is altered with age and in diseased states.

## Results

### Brain-wide calcium imaging captures a developmental shift in neural excitability

To understand how the intrinsic excitability of the vertebrate brain changes across early development, we performed calcium imaging using light-sheet microscopy. We visualized neural activity in live larval zebrafish at two developmental ages: 4 days post-fertilization (dpf), when larvae exhibit immature but functionally active neural circuits, and 9 dpf, a later stage reflecting more mature brain architecture and synaptic refinement. These stages were selected to capture developmental contrasts in baseline and stimulated neural activity. For this, we employed the *Tg*(HuC:GCaMP5G) transgenic line, which enables pan-neuronal monitoring of calcium activity and pentylenetetrazole (PTZ), a widely used chemo-neural stimulant that blocks GABA_A_ receptors to evoke neural hyperexcitation.

At 4 dpf, the recordings revealed robust and globally distributed patterns of spontaneous neural activity and large-amplitude, temporally structured calcium transients upon PTZ application (Supplementary Movie S1). Further analysis of raw calcium traces from a region of interest (ROI) in the dorsal forebrain (Fig. 1a) demonstrated clear, recurring periodic elevations in fluorescence with sharp rise times and slow decay kinetics (Fig. 1b). In stark contrast to the 4 dpf recordings, 9 dpf larvae displayed a different neural response to the same PTZ stimulus. Despite a significant increase in fluorescence intensity, the raw calcium traces lacked the rhythmic, large-amplitude events seen in 4 dpf larvae (Fig 1b’). Instead, activity was lower in amplitude and lacked the consistent bursting patterns that characterized the younger larvae (Supplementary Movie S2).

**Figure 1.**
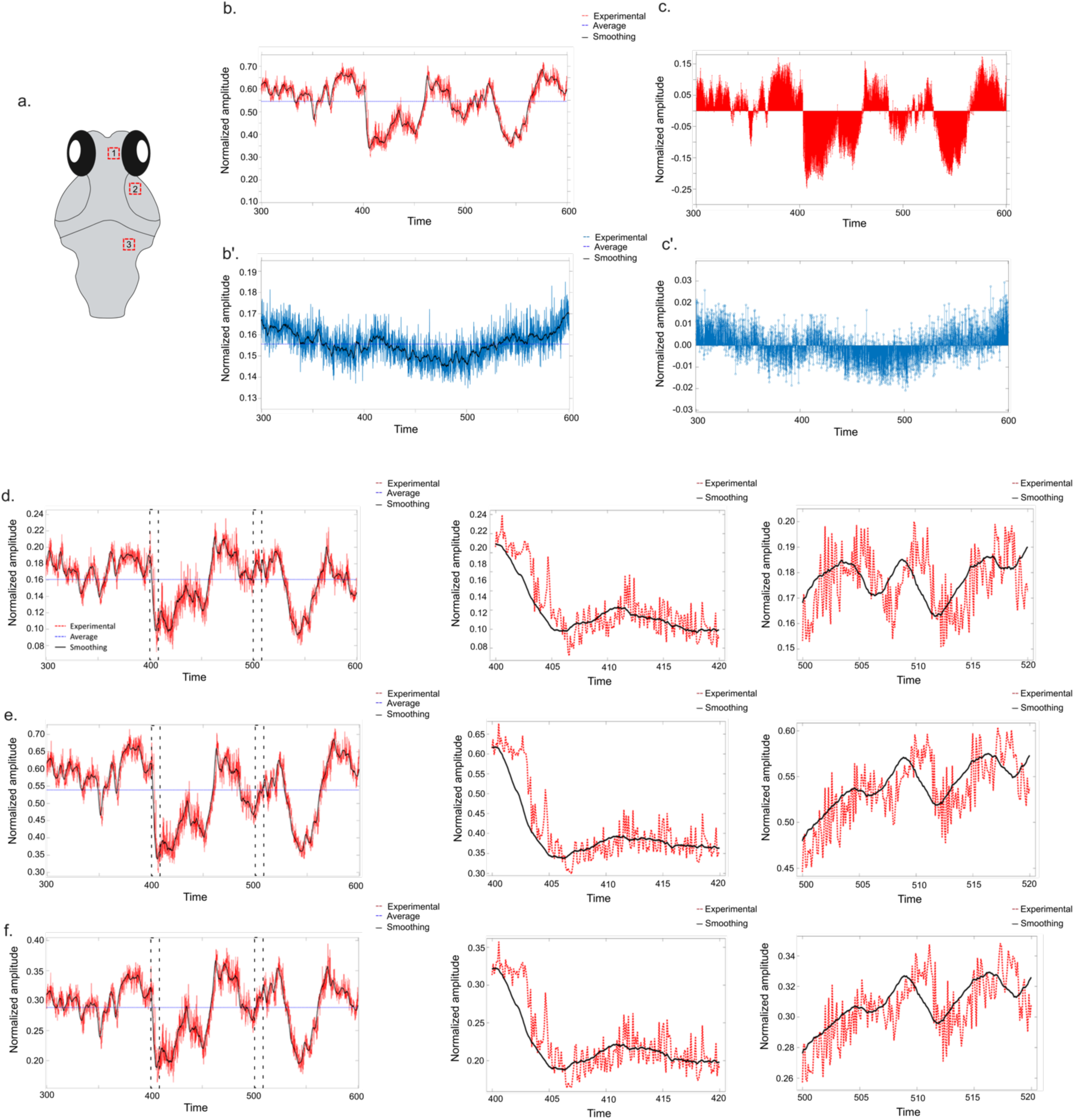
Developmental shift in neuronal excitability in larval zebrafish. **(a)** Schematic dorsal view of the zebrafish larval brain illustrating selected anatomical regions of interest (ROIs) used for calcium signal extraction and activity analysis. Red dashed boxes indicate the following regions: (1) telencephalon, (2) optic tectum, and (3) hindbrain. These ROIs were selected for investigation of neural excitability dynamics using single-plane images of fluorescence in the brains of live Tg(HuC:GCaMP5G) zebrafish larvae at 4 days post-fertilization (dpf), acquired using Mλ-sMx-SPIM light-sheet microscopy following exposure to 5 mM PTZ. **(b, b′)** Time-series plots of normalized calcium fluorescence (ΔF/F₀) traces from the 2^nd^ ROI from optic tectum in 4 dpf (red trace, b) and 9 dpf (blue trace, b′) larvae during the recording window (300–600 seconds following addition of PTZ). At 4 dpf (b), the signal shows prominent high-amplitude fluctuations, indicative of hyperexcitable neuronal activity, while at 9 dpf (b′), fluorescence signals remain near baseline with minimal transients, reflecting reduced excitability (note the difference in range of y axis). **(c, c′)** Deviation plots from the mean fluorescence signal (ΔF) at 4 dpf (c) and 9 dpf (c′) quantified from 2^nd^ ROI representing optic tectum. The larger deviations at 4 dpf (red trace, c) reflect burst-like and synchronous activity characteristic of a hyperexcitable neural network, whereas minimal deviation at 9 dpf (blue trace, c′) supports the phase of a more stable and less reactive network response to stimulus (note the difference in range of y axis). **(d, e, f)** Time-series analysis from selected ROIs corresponding to telencephalon (d), optic tectum (e), and hindbrain (f). Each ROI panel shows a full activity trace (300–600 s), along with two magnified 20-second segments extracted from the trace: t₁ (400–420 s): This time window captures a decline in both amplitude and frequency of calcium transients, representing an inter-burst interval, and further t₂ (500–520 s) plot window shows high-amplitude and frequent transients. The similar patterns in activity suggest synchronized patterns of excitability across the brain at 4 dpf. Red dashed lines represent raw signals; black lines indicate smoothed traces. The blue horizontal line denotes the mean fluorescence baseline used for normalization.

Computation of deviation-from-mean traces revealed substantial deviations from baseline following PTZ exposure in 4 dpf but not in 9 dpf larvae (Fig. 1c, c’). To assess if the hyperactivity observed in the 4 dpf brains was a general phenomenon, we analyzed multiple ROIs across anatomically distinct brain regions, including the telencephalon, midbrain, and cerebellar regions and observed parallel, high-amplitude activity consistently across regions (Fig. 1d-1f). These dynamics resembled global bursts or "network events", consistent with previously described patterns of network synchronization during immature circuit states [12].

Taken together, these findings reveal a clear shift in brain-wide responsiveness to excitatory stimulation with age, suggesting that early neural circuits undergo a transition in excitability during development that can potentially render the developing brain vulnerable to perturbations.

### Large locomotor events provide a behavioural readout of brain excitability

In order to establish an objective, quantifiable and reliable measure of neural excitability in larval zebrafish, we quantified PTZ-induced behavioural changes in 6 dpf larvae, which show robust and quantifiable locomotor behaviour. We conducted a systematic dose-response analysis using PTZ concentrations ranging from 0.5 mM to 5 mM. covering a broad dynamic range from subthreshold to near-saturating and analyzed the locomotor behaviour following treatment. The locomotor traces illustrate a concentration-dependent escalation in activity dynamics following PTZ exposure (Fig. 2a), capturing the emergence of rapid, large-amplitude motor bouts consistent with heightened excitability.

**Figure 2.**
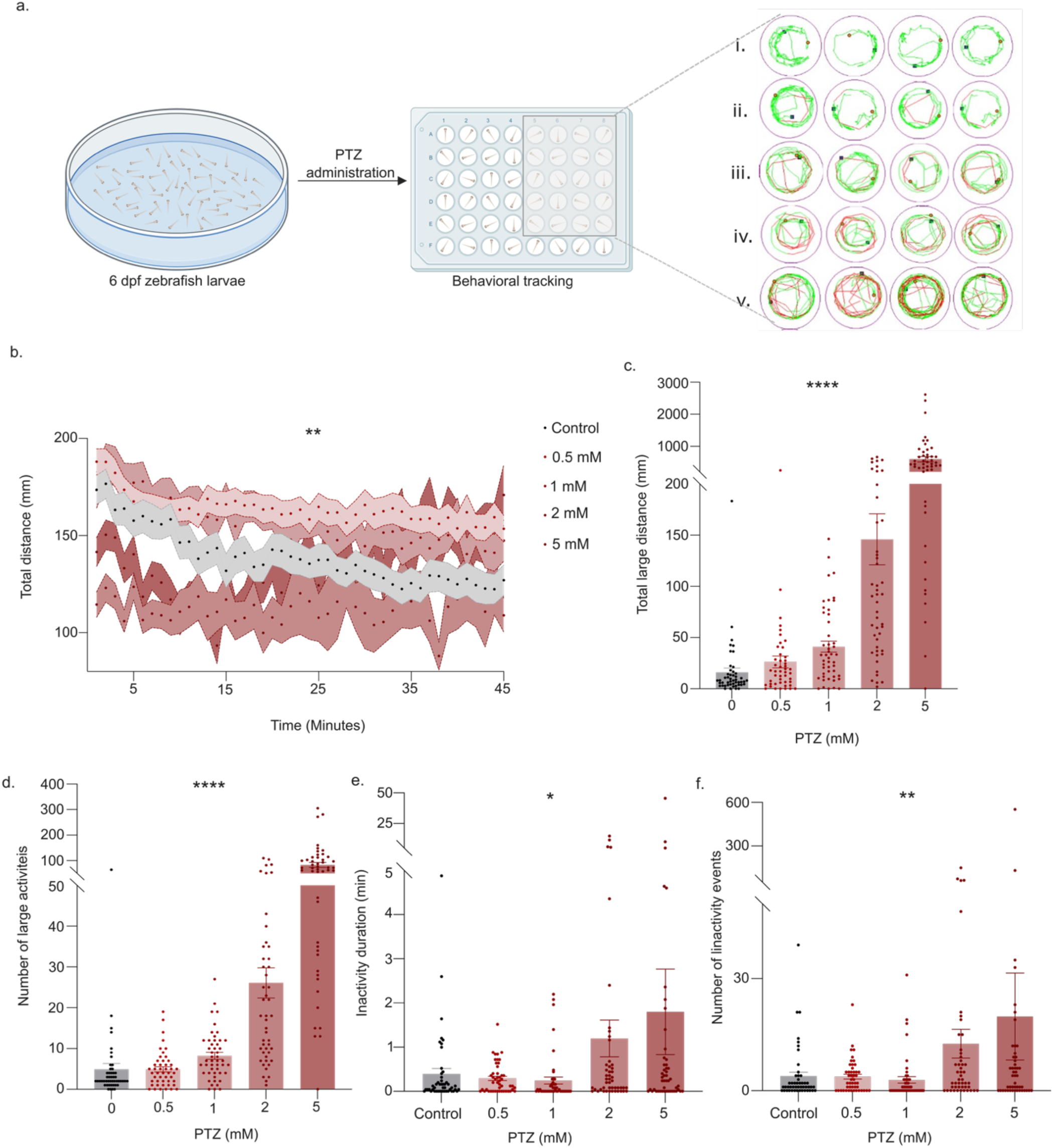
Specific large activity events provide a reliable means to quantify neural excitability. **(a)** Schematic representation of behavioral tracking of PTZ-induced activity in zebrafish larvae. Individual larvae were individually transferred into wells of a 48-well plate and behavior was recorded post-treatment. Representative one minute bin swim trajectories of individual larvae at increasing PTZ concentrations, visualized using Zebralab (Viewpoint, France). (i) Control (untreated), (ii) 0.5 mM PTZ, (iii) 1 mM PTZ, (iv) 2 mM PTZ, and (v) 5 mM PTZ. Larval locomotion in each well is color-coded: green for baseline locomotion (<45 mm/sec), red for hyperactivity (>45 mm/sec), and white for inactivity (0 mm/sec). Blue squares indicate the starting position, red dots denote the endpoint. As PTZ concentration increases, larvae exhibit hyperactive locomotory bursts and collapse (see Supplementary Movie S3). **(b)** Total distance (mm) swum per 1-minute time bin over 45 minutes during exposure to PTZ. Two-way repeated measures ANOVA showed a significant interaction between treatment and time (F(176, 10530) = 1.318, p =0.0033) and a significant effect of treatment (F(4, 10530) = 185, p < 0.0001); the main effect of time was significant (F(44, 10530) = 2.110; p < 0.0001). **(c)** Total large distance (mm) traveled as quantified from high-speed bursts of activity (>45 mm/s). Kruskal-Wallis test (p < 0.0001) with Dunn’s post hoc test: control vs. 0.5 mM (ns, p = 0.727), control vs. 1 mM (p = 0.0041), control vs. 2 mM ( p < 0.0001), control vs. 5 mM ( p < 0.0001). **(d)** No. of large or high-speed (>45 mm/s) events. Kruskal-Wallis test (p < 0.0001) with Dunn’s post hoc test: control vs. 0.5 mM (p > 0.9999), control vs. 1 mM (p = 0.0083), control vs. 2 mM (p < 0.0001), control vs. 5 mM (p < 0.0001). **(e)** Total inactivity duration (seconds). Kruskal-Wallis test (p = 0.0104); post hoc Dunn’s test: control vs. 0.5 mM (p = 0.5094), control vs. 1 mM (p = 0.1096), control vs. 2 mM (p = 0.1065), control vs. 5 mM (p = 0.1793). **(f)** Number of episodes of inactivity (0 mm/s). Kruskal-Wallis test (p = 0.0064); post hoc Dunn’s test: control vs. 0.5 mM (p > 0.9999), control vs. 1 mM (p = 0.7760), control vs. 2 mM (p = 0.1342), control vs. 5 mM (p = 0.5391). Sample size: n > 45 larvae per group. Data represent mean ± SEM. ns = not significant; ****p < 0.0001; ***p ≤ 0.001; **p < 0.01; *p < 0.05.

PTZ-exposed larvae displayed distinct changes in locomotor dynamics consistent with heightened excitability, including episodes of abrupt hyperactivity, rapid “whirlpool” movement bursts (≤1–3 sec), and intermittent inactivity (Supplementary Movie S3). While traditional metrics such as overall locomotor distance did reveal general increases in activity following PTZ treatment, the measure was not uniformly continuous across all time points or doses (Fig. 2b). Upon closer observation, we found that total locomotion integrates both baseline activity and seizure-associated hyperactivity, masking specific details of excitability-driven bursts behaviour. In contrast, when we defined and extracted "large locomotor events" as high-speed swimming bouts exceeding 45 mm/s, a threshold empirically derived from baseline recordings of control larvae which demonstrated such events in <1% of frames, PTZ-treated larvae consistently exceeded this threshold during episodes of heightened excitability.

Analysis of large locomotor events revealed a clear dose-dependent increase in high-speed (>45 mm/s) distance (Fig. 2c) and the frequency of such events (Fig. 2d) with increasing PTZ concentrations. Notably, concentrations of 1 mM and above elicited robust, statistically significant increases in large activity, while 2 mM and 5 mM PTZ induced near-maximal responses. These results establish large movement events as a specific and reliable behavioral correlate of neural excitability than gross locomotor activity alone. In addition to hyperactivity, we also characterized behavioral pauses (speed = 0 mm/s), which tended to increase in both duration (Fig. 2e) and frequency (Fig. 2f) at higher PTZ doses. These episodes typically followed intense bursts of activity and were often marked by abrupt behavioral transitions, including sudden freezing, loss of posture, or tight circular swimming, suggesting rapid shifts in excitability state.

Together, our data demonstrates that large locomotor events serve as a robust, sensitive, and physiologically relevant behavioral proxy for brain excitability in larval zebrafish.

### A critical developmental window of reduced neural excitability

To determine whether the intrinsic changes in brain excitability observed through live imaging also manifest at the behavioral level, we performed longitudinal behavioral profiling of naïve zebrafish larvae subjected to (PTZ) treatment (Fig. 3a). Zebrafish larvae from a single clutch were tracked every day from 4 to 13 days post-fertilization (dpf) to map developmental changes in behavioral responsiveness to PTZ-induced excitability. In untreated controls, baseline swimming gradually increased with age and plateaued around ∼8 dpf (Fig. 3b). In contrast, larvae exposed to PTZ displayed a non-linear trajectory of excitability-associated behavior. From 4 to 7 dpf, PTZ-treated larvae exhibited pronounced sensitivity, with large movements peaking at 6 dpf, indicating a phase of maximal network excitability (Fig. 3c, d). This heightened excitability declined by 9 dpf, with a pronounced drop in both total large-movement distance and the frequency of hyperactive events (Fig. 3c, d, e; Supplementary Movie S4). Overall, this temporal shift in excitability was consistent across PTZ concentrations, as similar trends were observed in both 2 mM and 5 mM PTZ-treated groups and in a separate wild-type zebrafish strain (Fig. S1).

**Figure 3.**
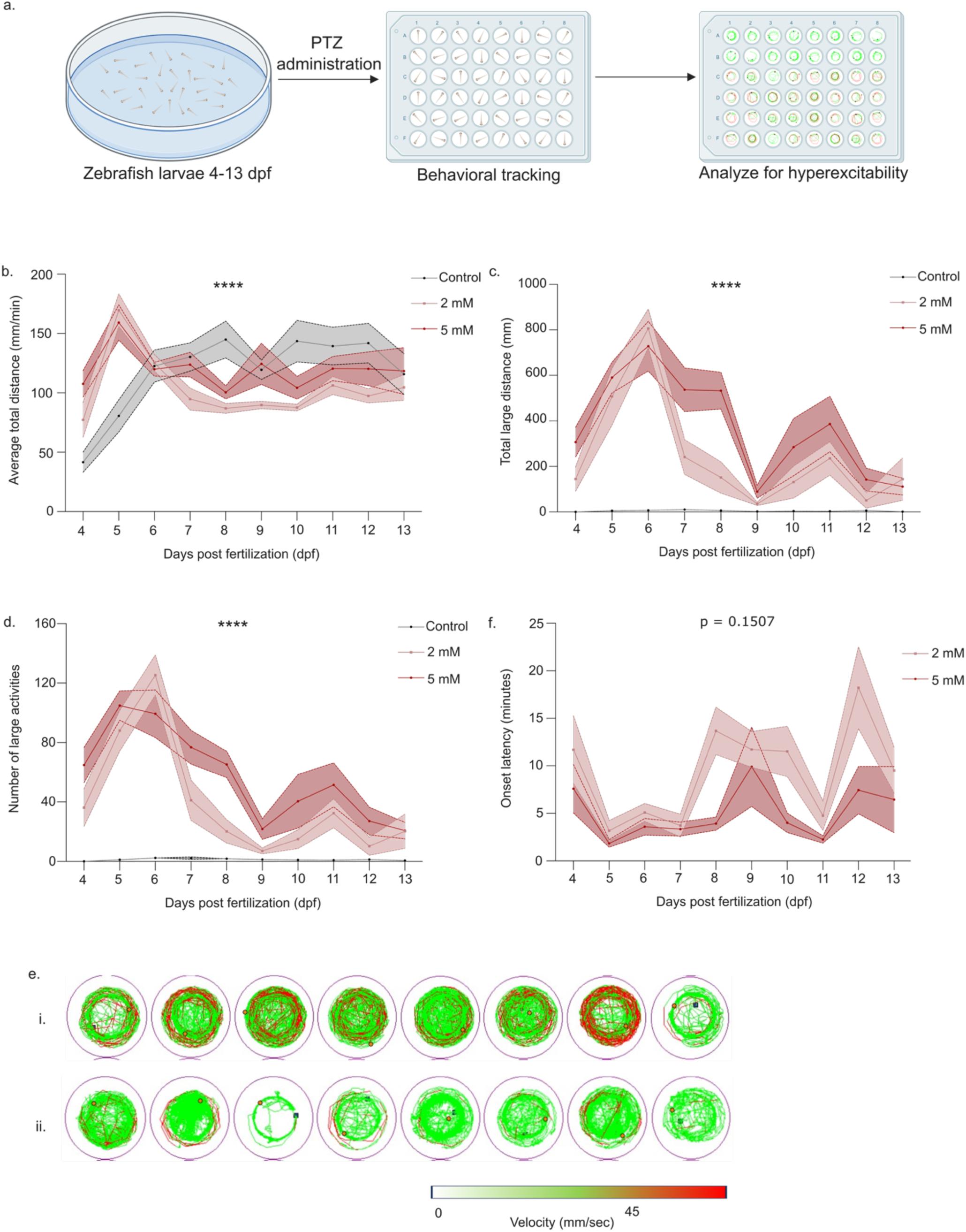
Behavioral response reveals a critical window of transition in neural excitability. **(a)** Experimental schematic for PTZ-responsive behavioral assessment across developmental stages. Zebrafish larvae from single clutch aged 4–13 days post-fertilization (dpf) were transferred to a 48-well plate, where they were exposed to pentylenetetrazole (PTZ) to induce hyperexcitable behavior. Following drug administration, larval behavior was recorded and analyzed to assess behavioral responses. **(b)** Average total distance swum (mm/min) by control and PTZ-treated larvae (2 mM, 5 mM) from 4 to 13 dpf. Two-way mixed-effects ANOVA, effect of age (F(6.152, 198.9) = 5.239, p < 0.0001); treatment (F(2, 33) = 3.826, p = 0.032), and interaction (F(18, 291) = 4.769, p < 0.0001). Post hoc Tukey’s test: 6 vs 9 dpf in control (p >0.9999), 6 vs 9 dpf in 2 mM PTZ group (p = 0.0058), 6 vs 9 dpf in 5 mM PTZ group (p >0.9999). **(c)** Large distance traveled during high-speed (>45 mm/s) events. Two-way mixed-effects ANOVA, effect of age (F(5.648, 182.6) = 15.34, p < 0.0001); treatment (F(2, 33) = 60.05, p < 0.0001), and interaction (F(18, 291) = 4.924, p < 0.0001). Post hoc Tukey’s test: 6 vs 9 dpf in control (p = 0.7817), 6 vs 9 dpf in 2 mM PTZ group (p <0.0001), 6 vs 9 dpf in 5 mM PTZ group (p = 0.0071). **(d)** Total number of high-speed (>45 mm/s) seizure-associated or large events. Two-way mixed-effects ANOVA, effect of age (F(6.255, 202.3) = 17.99, p < 0.0001); treatment (F(2, 33) = 82.42, p < 0.0001), and interaction (F(18, 291) = 5.580, p < 0.0001). Post hoc Tukey’s test: 6 vs 9 dpf in control (p = 0.9761), 6 vs 9 dpf in 2 mM PTZ group (p <0.0001), 6 vs 9 dpf in 5 mM PTZ group (p = 0.0611). **(e)** Representative traces of locomotion of (i) 6 and (ii) 9 dpf larvae; baseline locomotion (green), seizure-like hyperactivity (red), and inactivity (white). Blue square indicates the starting position; red dot marks the endpoint. **(f)** Latency (in minutes) to the first high-speed event post-PTZ exposure. Two-way mixed-effects ANOVA, effect of age (F(4.495, 91.91) = 5.362, p = 0.0004); treatment (F(1, 22) = 19.87, p = 0.0002), and interaction (F(9, 184) = 1.500, p = 0.1507). Post hoc Tukey’s test: 6 vs 9 dpf in 2 mM PTZ group (p = 0.0728), 6 vs 9 dpf in 5 mM PTZ group (p = 0.8779). Sample size: n = 12 larvae per group per day. Data represent mean ± SEM. ns = not significant; ****p < 0.0001; ***p ≤ 0.001; **p < 0.01; *p < 0.05.

A partial resurgence of large locomotor events was observed from 10 to 13 dpf, although activity levels remained consistently lower than the early high-excitability phase (Fig. 3c, d). Control larvae across all time points displayed negligible large-movement activity, confirming that the observed dynamics were stimulus-specific and not attributable to spontaneous developmental shifts in baseline motor behavior.

Latency to the first large locomotor event following PTZ exposure was shortest during the early phase (4–7 dpf), reflecting basal hyperexcitability, and peaked at 8–10 dpf (Fig. 3f). Taken together, these data overlap with our observations from the brain activity mapping and define 8–9 dpf as a critical transition phase during which the zebrafish brain shifts from a hyperexcitable to a relatively resistant state.

Further, to examine whether repeated neural excitation alters the intrinsic developmental pattern of brain excitability, we subjected larval zebrafish to daily treatment with PTZ across a 10-day window (4–13 dpf). 1 mM and 2 mM PTZ were used since chronic exposure to high PTZ concentrations (2 mM and 5 mM) resulted in substantial mortality (Fig. S2). Across the 10-day window, large locomotor activity (>45 mm/s) exhibited a progressive decline following the first exposure at 4 dpf. This decline suppressed the characteristic early excitability peak typically observed at 5–6 dpf in untreated animals (Fig. S3b, c). Interestingly, during the 8–9 dpf window, when naïve larvae exhibit a pronounced reduction in excitability, chronically stimulated larvae displayed a paradoxical increase in high-speed activity. This was also evident with the transient reduction in latency at 8–9 dpf (Fig. S3d). The PTZ-treated group showed deviation in average total distance (mm/min) compared to the control, yet the overall pattern across time remained similar, reflecting normal development of locomotion (Fig. S3e). These results suggest that chronic neural stimulation perturbs the brain’s intrinsic developmental programming and its homeostatic mechanisms.

### Molecular markers are altered across developmental stages mirroring brain excitability

To investigate whether the temporally regulated behavioral and neural excitability patterns observed during zebrafish development are reflected at the molecular level, we examined two widely used markers of neuronal activation: *c-fos*, an immediate-early gene responsive to neuronal activity, and the ratio of phosphorylated (pERK; phosphorylated extracellular signal-regulated kinase) to total ERK (tERK; total extracellular signal-regulated kinase), which represents rapid signaling events downstream of synaptic activity.

At 6 dpf, when larvae show hyperexcitability, PTZ exposure resulted in a significant upregulation of *c-fos* compared to untreated controls, while this transcriptional response was markedly attenuated at 9 dpf, reflecting a blunted excitability response (Fig. 4a). A modest re-emergence of *c-fos* induction was observed by 13 dpf, consistent with the partial behavioral rebound seen during this stage. Assessment of pERK/tERK ratios showed a robust increase in the ratio in 6 dpf larvae indicating elevated synaptic activity in response to PTZ (Fig. 4b, c). However, no statistically significant induction of pERK/tERK was detected at 9 dpf. These molecular signatures of neural activation closely parallel the functional transitions observed via light-sheet imaging and behavioral assays, reinforcing the presence of a defined developmental window during which the brain’s excitability heightened.

**Figure 4.**
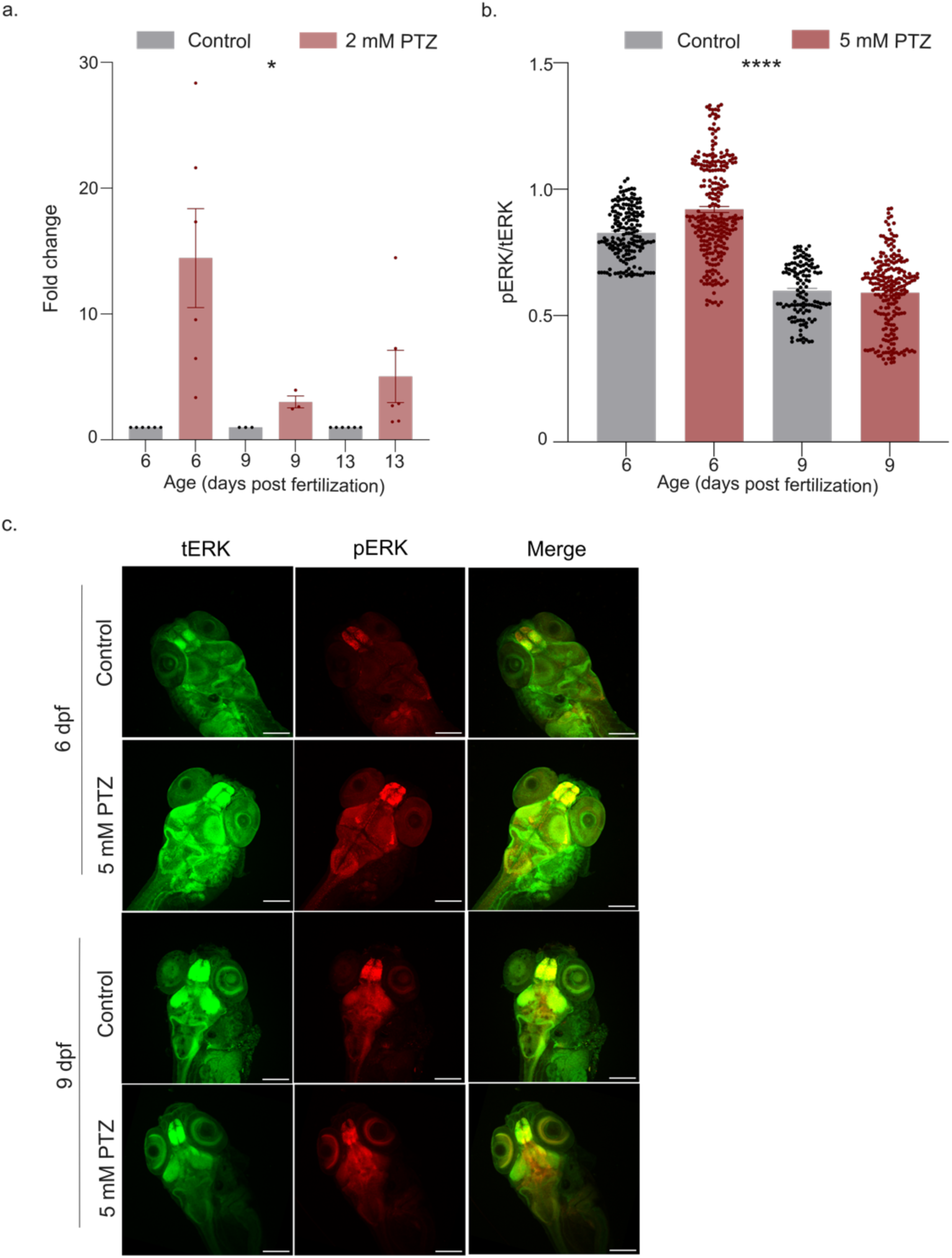
Molecular readouts reflect age-dependent alterations in excitability. **(a)** RT-qPCR analysis of c-fos transcript levels in naïve larvae exposed to 2 mM PTZ at 6 dpf (n = 6), 9 dpf (n = 3), and 13 dpf (n = 6). Two-way mixed-effects ANOVA, effect of age, (F(2, 24) = 3.776, p = 0.0375); treatment (F(1, 24) = 11.53, p = 0.0024), interaction (F(2, 24) = 3.776, p = 0.0375). Post hoc Tukey’s test: control vs 2mM PTZ (p = 0.0012) at 6 dpf; control vs 2mM PTZ (p = 0.9958) at 9 dpf. **(b)** Quantification of the pERK/tERK fluorescence intensities in the telencephalon of 6 dpf (n = 10) and 9 dpf (n = 8) larvae treated with 5 mM PTZ. Age-matched controls: 6 dpf (n = 6,), 9 dpf (n = 3). Approximate 25-30 slices per n. Kruskal-Wallis test (p < 0.0001) with Dunn’s post hoc test: control vs. 5 mM (p = 0.0302) at 6 dpf, and control vs. 5 mM (p > 0.9999) at 9 dpf. **(c)** Representative whole-mount immunofluorescence images of the brain showing total ERK (tERK; green), phosphorylated ERK (pERK; red), and merged channels (co-localization; yellow) in naïve 6 and 9 dpf larvae treated with 5 mM PTZ, along with respective age-matched controls. Scale bars: 200µm.

Put together, using a combination of behavioural and molecular readouts, we demonstrate that the developing vertebrate brain shows clear phases of hyperexcitability when it is susceptible to external perturbations and this has potential implications in understanding the etiology of neurodevelopmental disorders.

## Discussion

The developing brain is particularly susceptible to internal and external perturbations, many of which can result in long-lasting consequences [2,3]. This heightened vulnerability is especially evident in early-onset neurological conditions such as epilepsy, where seizures are most likely to emerge during infancy and early childhood [13-17]. Though an understanding of how intrinsic brain excitability fluctuates during this period is vital, gaining insight into these processes has been limited by the challenges of restricted real-time access to mammalian embryos.

In this study, we employed the widely used GABA_A_ receptor antagonist pentylenetetrazole (PTZ) as a stimulant for evoking excitation [18]. We then combined light-sheet calcium imaging, behavioral assays and molecular outputs to analyze the developmental dynamics of neural excitability in zebrafish larvae. Our longitudinal approach revealed a previously underrecognized developmental shift in brain excitability between 4 and 9 dpf, a time frame that corresponds to early postnatal stages in rodents and human infants [19].

At 4 dpf, PTZ evoked a robust and globally synchronized response across the larval brain, while at 9 dpf, the same PTZ treatment produced a comparatively muted and less synchronized activation pattern. This biphasic excitability profile aligns well with observations from rodent seizure models, where the early postnatal brain is markedly more susceptible to convulsants as reported in rats [7,20,21,22,23] and mice [24]. The reduced *c-fos* and pERK activation observed at 9 dpf may indicate decreased synaptic gain, more efficient inhibitory control, or other circuit-level changes in excitability regulation.

A particularly notable observation was the modulation of neural excitability by experience. Repeated PTZ exposure led to a suppression of the early hyperexcitable state seen at 4–6 dpf and induced a resurgence of sensitivity at 9 dpf, a time when naïve larvae were otherwise resistant. This phenomenon stands in contrast to traditional kindling models in rodents, where repeated stimulation progressively lowers seizure thresholds and amplifies seizure severity over time [7]. Our results suggest a different mechanism may be at play in zebrafish, where early and repeated convulsant exposure appears to disrupt the normal trajectory of excitability maturation.

The prevailing hypothesis for heightened early-life excitability is the developmental trajectory of chloride transporter expression, particularly those of the SLC12 family. The ontogenetic switch from NKCC1, which imports chloride and renders GABA depolarizing, to KCC2, which exports chloride and enables GABAergic inhibition, is known to underlie the shift in GABAergic signaling polarity from excitatory to inhibitory during brain maturation [25,26]. Although *kcc2* is expressed as early as 2 dpf in zebrafish, the precise temporal dynamics and regional expression patterns of NKCC1 and KCC2 remain under-characterized [27]. Their ratio may modulate GABA polarity and thereby contribute to the observed changes in PTZ sensitivity across developmental stages. It is also possible that other neurodevelopmental processes such as synaptogenesis, axonal pruning, myelination, or neurotransmitter receptor composition contribute to the excitability landscape during development.

Interestingly, earlier studies investigating the basis of visually guided behavior have demonstrated that spontaneous neural activity in zebrafish larvae undergoes significant developmental reorganization around 5–6 dpf. From 8 dpf onwards, this activity reflects a period of circuit stabilization following a phase of heightened plasticity and network refinement [28]. These observations which suggest a developmental shift are in line with our observations, indicating that the changes we observe are shaped by the maturation of the circuitry involved.

These findings carry important implications for epilepsy research, especially because it is well-established that early-life seizures can remodel neural circuits and increase the risk of later-life cognitive deficits [29]. First, they reinforce the idea that seizure susceptibility is developmentally programmed and follows a non-linear trajectory, rather than remaining uniformly high or low across early life, which supports the observation of significantly higher incidence of seizure disorders during early childhood and in the elderly population. Second, since PTZ acts by antagonizing GABA_A_ receptors, the altered response observed at 9 dpf may model a developmental “checkpoint” or a “critical period”, and understanding such checkpoints may help in the design of age-specific diagnostic, prognostic, or therapeutic strategies.

While our study focused on a single pharmacological agent, PTZ, this approach has some limitations. Despite its well-characterized GABA_A_ receptor antagonist profile, PTZ alone may not capture the full spectrum of excitability responses. Future investigations could explore whether similar developmental patterns are observed with other excitatory agents, such as kainic acid, NMDA receptor agonists, or 4-aminopyridine, or in response to non-pharmacological stressors like hypoxia, metabolic imbalance, or sensory deprivation. Additionally, genetic manipulation of NKCC1, KCC2, or other GABAergic signaling components would provide causal insight into how inhibitory balance shapes excitability development. More detailed spatial and temporal analysis, including ROI-based synchrony measurements and region-specific gene expression profiling, could further refine our understanding of how excitability transitions manifest across brain regions and circuits.

In summary, our study reveals a clear previously underrecognized developmental shift in brain excitability during zebrafish development during a period that corresponds roughly to early postnatal stages in rodents and human infants. These findings will help in further studies on circuit homeostasis, neurodevelopmental processes, and age-specific risk to neurological diseases.

## Materials and Methods

### Fish husbandry and fish lines

Adult wildtype zebrafish (*Danio rerio*) and *Tg*(HuC:GCaMP5G) were maintained in mixed sex groups in a GenDanio zebrafish housing system under standard conditions of 28.5°C, pH 7, and conductivity 300 μS/cm under a 14 h/10 h light/dark cycle. Fish were bred and reared at 28.5°C according to standard protocols. Embryos were maintained at optimal density and incubated at 28.5°C in system water supplemented with 0.01 mg/L methylene blue until 6 days post-fertilization (dpf). Larvae were transferred into larger containers at 6 dpf, and feeding was initiated thrice/day. Experimental procedures were approved by the Institutional Animal Ethics Committee of the Indian Institute of Science Education and Research, Thiruvananthapuram, India (IISERTVM/IAEC/2024-XI/FormB-03-R2). All procedures were in accordance with CCSEA guidelines.

### Sample Preparation and Embedding

For light-sheet recordings, non-anesthetized *Tg(HuC:GCaMP5G)* zebrafish larvae were embedded in 1.5% ultrapure low-melting-point agarose (ThermoFisher, #16520050), prepared in filtered fish facility water. Agarose was dissolved by heating (75–80 °C) and equilibrated to ∼37 °C in a dry bath to prevent thermal stress. A small volume of cooled agarose (∼33 °C) was added in a petri dish, and each larva was aspirated with a 200 μL micropipette along with a small plug of agarose into glass capillaries (Sigma-Aldrich, #Z114995-100EA). A thin pin was inserted into the capillary to displace some agarose and create a small chamber at the anterior end for drug addition during imaging and let to stay till the agarose solidified. To prepare the sample for imaging, the agarose cylinder containing the larva was gently extruded from the capillary tip from the anterior end, ensuring that the head protruded out of the capillary. The posterior end was sealed with additional agarose and polytetrafluoroethylene (PTFE) film. Larvae were maintained at room temperature (∼22–25 °C) throughout the procedure.

### Light-Sheet Imaging and Data Analysis

Multi-wavelength Simultaneous Multiple-level Magnification Selective Plane Illumination Microscopy (Mλ-sMx-SPIM) system as previously described [30], was used for calcium imaging. The system uses a tunable laser source for the illumination and dual detection arms for the image recording. For this study, the wavelength was tuned to 488 nm and a detection arm equipped with a 4× objective lens (TL4X-SAP) and a tube lens (f = 100mm), hence a total of 4.44× magnification. A long-pass filter (FEL0500, Thorlabs, Inc., *λ_cut-on_* = 500 nm) was placed before the camera to remove the excitation beam. Capillary tubes were fixed on a custom 3D-printed stage and mounted on the sample mounting stage of the imaging system. A single imaging plane within the brain was selected and recorded continuously at 10 frames per second using a Blackfly S USB3 camera (model: BFS-U3-32S4M-C). Each recording session began with a 2-minute baseline acquisition in system water. Following baseline acquisition, pentylenetetrazole (PTZ; Sigma-Aldrich, P6500) was added to the chamber to a final concentration of 5 mM, without disturbing the capillary-embedded larva. Imaging continued for an additional 8 minutes, capturing the spatiotemporal dynamics of PTZ-induced excitability *in vivo*.

The acquired time-lapse image stacks were processed using custom MATLAB scripts. Regions of interest (ROIs) were selected based on anatomical landmarks (e.g., telencephalon, tectum, hindbrain), and fluorescence traces (ΔF/F₀) were extracted, where F₀ is the maximum fluorescence intensity. Traces were smoothed using a moving average filter (averaging over 25 frames, i.e., 250 milliseconds) to visualize the nature of fluorescence intensity variations and highlight oscillatory patterns. Deviation-from-mean analyses and trace segment extractions were performed to assess signal periodicity, event amplitude, and dynamic changes in activity over time.

### Drug Treatment and Behavior Recording

Pentylenetetrazole (PTZ; Sigma-Aldrich, #P6500) was dissolved in water to prepare 5× stock solutions, which were diluted to final concentrations of 0.5 mM, 1 mM, 2 mM, and 5 mM, with control groups receiving system water only. Larvae were individually transferred to wells, acclimated for 10 minutes to minimize stress, and recorded immediately after drug administration using a ZebraBox™ automated tracking system for 45 minutes (Viewpoint, Lyon, France). Recordings were performed under constant 1040 lux lighting, between 9:00 AM and 4:00 PM to control for circadian variability. Behavioral data were analyzed using ZebraLab™ software (Viewpoint, Lyon, France), with locomotor states classified as baseline activity (<45 mm/s), seizure-associated hyperactivity (≥45 mm/s), and inactivity (0 mm/s).

### Real-time quantitative polymerase chain reaction

Total RNA (300 ng) was reverse-transcribed into cDNA using the PrimeScript RT Master Mix (Perfect RealTime; Takara Bio, #RR036A) according to the manufacturer’s protocol. qPCR reactions were performed using SYBR Green Master Mix (Invitrogen) on a Light Cycler Bio-Rad (CFX connect) (primer details in Supplementary Information). *β-actin* gene (*actb1*) was used as the reference gene for normalization, and relative gene expression was calculated using the ΔΔCt method. Each *n* represents pooled RNA from 3 larvae

### Whole Mount Immunostaining

Zebrafish larvae were reared in system water supplemented with 0.003% 1-phenyl-2-thiourea (PTU) beginning at 10 hours-post fertilization. Following PTZ treatment, larvae were immediately fixed in 4% paraformaldehyde (PFA) in phosphate-buffered saline (PBS) at 4°C for 24 hours. Fixed larvae were washed thrice in 1xPBS with 0.1% Tween 20 (PBT) at room temperature (RT). Antigen retrieval was performed by incubating larvae in 150 mM Tris-HCl (pH 9.0) in PBT for 5 minutes, followed by incubation at 70°C for 15 minutes, followed by three PBT washes. Larvae were permeabilized in 10 µg/mL proteinase K in PBT for 20 minutes for 6 dpf and 22 minutes for 9 dpf larvae, washed three times in PBT, and blocked in blocking solution (5% fetal bovine serum in PBT) for ≥1 h at RT. Primary antibodies against phosphorylated ERK (pERK; Cell Signaling Technology, #4370T) and total ERK (tERK; Cell Signaling Technology, #4696T) were diluted 1:500 in blocking solution and applied to larvae overnight at 4°C. After three PBT washes, larvae were incubated with Alexa Fluor 561- and 488-conjugated secondary antibodies (Invitrogen) diluted 1:300 in blocking solution overnight at 4°C. Following three final PBT washes (15 min each), fluorescence signals were verified using a fluorescence microscope. Larvae were mounted in 75% glycerol in PBT and imaged with Olympus fluoview FV3000 inverted microscope using a 10× objective.

### Statistical Analysis

Data are presented as the mean ± standard error of the mean (SEM). Statistical analyses were conducted using GraphPad Prism version 8.0.1 (244) software. Depending on the dataset and experimental design, statistical significance was calculated using the Kruskal–Wallis test, two-way mixed-effects ANOVA, or two-way repeated measures ANOVA, as applicable. Post-hoc analyses included Dunn’s or Tukey’s multiple comparisons tests, where appropriate. Statistical differences were indicated as * *p* ≤ 0.05, ** *p* ≤ 0.01, *** *p* ≤ 0.001, **** *p* ≤ 0.0001, with exact values provided in figure legends where appropriate.

## Acknowledgements

This work was supported by the Indian Institute of Science Education and Research Thiruvananthapuram and grants from the Department of Biotechnology (BT/PR52416/BMS/85/208/2024) and Anusandhan National Research Foundation (ANRF/ECRG/2024/004451/LS). AS is a Rising Stars awardee from the International Brain Research Organization. GS is supported by a fellowship from the University Grants Commission, India. We are grateful to the support extended by the microscopy facility and core instrumentation facility at the School of Biology, IISER Thiruvananthapuram. We thank Ms. Anagha Muralidharan and Ms. Malavika KA for their assistance with standardizing sample preparation protocols for the light sheet microscopy.

## Author contributions

Conceptualization: PS and AS; Methodology: PS, GS, AKS; Investigation: PS, GS, AKS, AP; Visualization: PS, GS, AKS, AP; Supervision: MSS, AS; Writing—original draft: PS, GS, AS; Writing—review & editing: PS, GS, MSS, AS

## Data and materials availability

All data are available in the main text or the supplementary materials.

