## Supplementary information for "A developmental transition from early neural hyperexcitability in zebrafish"

### A. Supporting figures

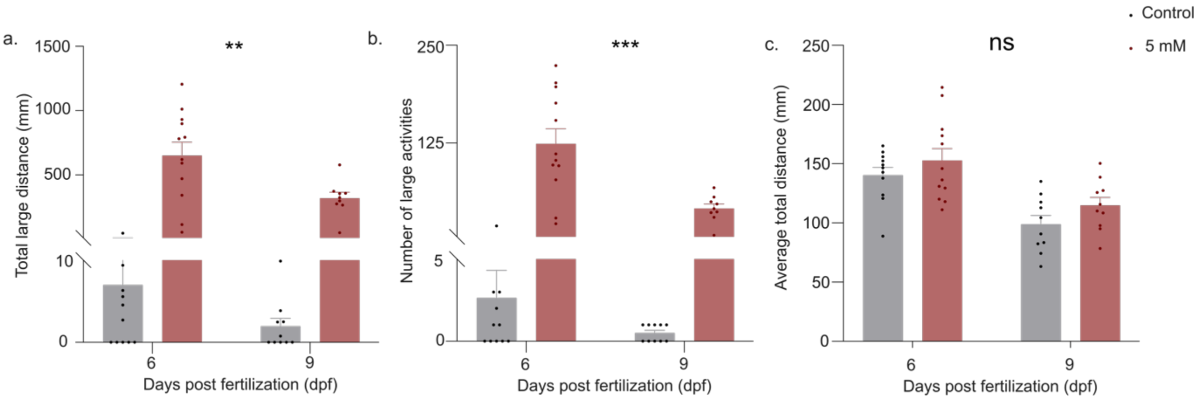


***Figure S1:*** *PTZ induced seizure behaviors in leopard wild-type zebrafish larvae at 6 and 9 dpf.*

***(a)*** *Large distance traveled during high-speed (>45 mm/s) events. Two-way ANOVA, effect of age (F(1, 38) = 7.248, p < 0.0105); treatment (F(1, 38) = 59.33, p < 0.0001), and interaction (F(1,38) = 6.822, p = 0.0128). Post hoc Sidak's test: 6 vs 9 dpf in control (p = 0.9979) and, 6 vs 9 dpf in 5 mM PTZ group (p = 0.0012).*

***(b)*** *Total number of high-speed (>45 mm/s) seizure-associated or large events. Two-way ANOVA, effect of age (F(1, 38) = 13.83, p < 0.0006); treatment (F(1, 38) = 50.54, p < 0.0001), and interaction (F(1,38) = 12.47, p = 0.0011). Post hoc Sidak's test: 6 vs 9 dpf in control (p = 0.9890) and, 6 vs 9 dpf in 5 mM PTZ group (p = <0.0001).*

***(c)*** *Average total distance swum (mm/min) by control and 5mM PTZ-treated larvae at 6 and 9 dpf. Two-way ANOVA, effect of age (F(1, 39) = 23.92, p < 0.0001); treatment (F(1, 39) = 3.074, p = 0.0874), and interaction (F(1,39) = 0.04332, p = 0.8362). Post hoc Sidak's test: 6 vs 9 dpf in control (p = 0.0019) and, 6 vs 9 dpf in 5 mM PTZ group (p = 0.0037).*

*Data represent mean ± SEM (n = 12 larvae/group). ns = not significant;* $****p < 0.0001$; $***p \leq 0.001$; $**p < 0.01$; $*p < 0.05$

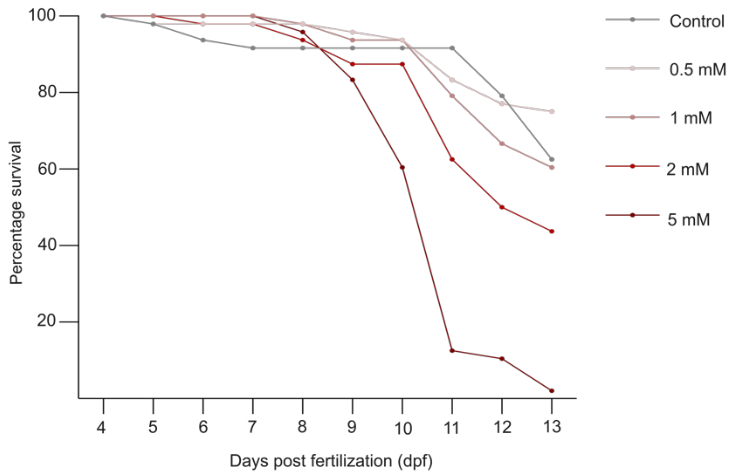


***Figure S2:*** *Survival rates of larvae exposed to 500 µM, 1 mM, 2 mM, and 5 mM PTZ from 4–13 dpf demonstrates an increase in dose-dependent lethality upon chronic PTZ treatment (n = 48/group).*

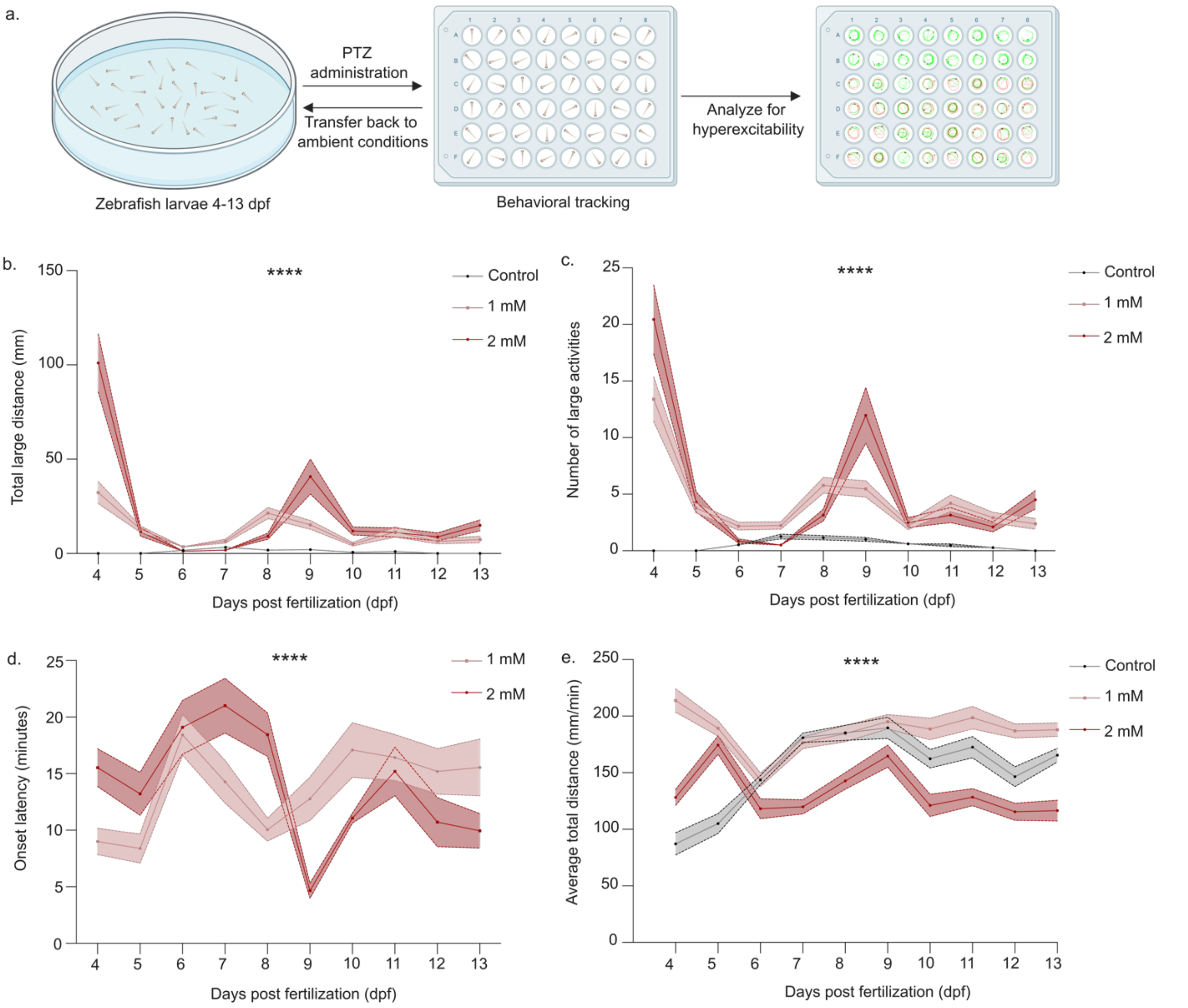


***Figure S3: Repeated PTZ excitation modulates the temporal intrinsic excitability in zebrafish larvae***

***(a)*** *Zebrafish larvae (4–13 days post-fertilization, dpf) were exposed daily to pentylenetetrazole (PTZ) for 45 minutes in a 48-well plate to induce neural hyperexcitability. Following each exposure, the larvae were transferred back to drug-free ambient conditions. The same cohort of larvae was subjected to this protocol across multiple developmental days, enabling daily behavioral assessment to evaluate cumulative effects on repeated excitability.*

***(b)*** *Total large distance traveled during high-speed (>45 mm/s) seizure-associated locomotion ("large events"). Chronic PTZ exposure attenuated peak sensitivity at 5–6 dpf. Two-way mixed-effects ANOVA, effect of age ($F(1.713, 201.0) = 28.51$, $p < 0.0001$); treatment ($F(2, 1056) = 52.56$, $p < 0.0001$), interaction ($F(18, 1056) = 16.60$, $p < 0.0001$). Post hoc Tukey's test: 4 vs 6 dpf ($p = 0.0003$), 6 vs 9 dpf ($p = 0.0030$) in 1 mM PTZ group; 4 vs 6 dpf ($p <0.0001$), 6 vs 9 dpf (0.0139) in 2 mM PTZ group.*

***(c)*** *Frequency of large events, showing transient resurgence at 8–9 dpf. Two-way mixed-effects ANOVA, effect of age ($F(2.261, 272.6) = 34.51$, $p < 0.0001$); treatment ($F(2, 1085) = 68.32$, $p < 0.0001$), interaction ($F(18, 1085) = 13.01$, $p < 0.0001$). Post hoc Tukey's test: 4 vs 6 dpf ($p < 0.0001$), 6 vs 9 dpf ($p = 0.0100$) in 1 mM PTZ group; 4 vs 6 dpf ($p <0.0001$), 6 vs 9 dpf (0.0039) in 2 mM PTZ group.*

***(d)*** *Latency (minutes) to the first high-speed event, with transient decrease at 8–9 dpf. Two-way mixed-effects ANOVA, effect of age ($F(7.236, 478.4) = 5.606$, $p < 0.0001$); treatment ($F(1, 595) = 0.04209$, $p = 0.8375$), interaction ($F(9, 595) = 6.044$, $p < 0.0001$). Post hoc Tukey's test: 4 vs 6 dpf ($p = 0.0067$), 6 vs 9 dpf ($p = 0.5761$) in 1 mM PTZ group; 4 vs 6 dpf ($p = 0.9729$), 6 vs 9 dpf (0.0005) in 2 mM PTZ group.*

***(e)*** *Total locomotion (mm) over time, reflecting normal motor development. Two-way mixed-effects ANOVA, effect of age , ($F(7.618, 906.5) = 9.623$, $p < 0.0001$); treatment ($F(2, 141) = 91.99$, $p < 0.0001$), interaction ($F(18, 1071) = 12.16$, $p < 0.0001$). Post hoc Tukey's test: 4 vs 6 dpf ($p = 0.0001$), 6 vs 9 dpf ($p = 0.0012$) in control; 4 vs 6 dpf ($p <0.0001$), 6 vs 9 dpf ($p <0.0001$) in 1 mM PTZ group; 4 vs 6 dpf ($p = 0.9963$), 6 vs 9 dpf (0.0262) in 2 mM PTZ group.*

*Sample size: $n > 45$ larvae/group at start of treatment. Data represent mean ± SEM. ns = not significant; ****$p < 0.0001$; ***$p \leq 0.001$; **$p < 0.01$; *$p < 0.05$.*

**B. List of supporting movies**

**Movie S1:** Representative real-time light-sheet imaging of PTZ-induced brain-wide excitability dynamics in 4 dpf zebrafish larvae

**Movie S2:** Representative real-time light-sheet imaging of PTZ-induced brain-wide excitability dynamics in 9 dpf zebrafish larvae

**Movie S3:** Representative video of PTZ-evoked behavioral hyperactivity in 6 dpf zebrafish larvae.

**Movie S4:** Altered PTZ-evoked behavioral response in 9 dpf zebrafish larvae compared to 6 dpf.

**C. List of primers used for qRT-PCR experiments**

| Sr. | Gene | Primer Sequence (5' → 3') | Amplicon (bp) |
|---|---|---|---|
| 1. | *c-fos*_FP | GGCGCAGCTCAATCCTACAA | 152 |
| 2. | *c-fos*_RP | GCAGCAGCCATCTTGTTTCG | |
| 3. | *β-actin*_FP | TCTTCACTCCCCTTGTTCACAA | 181 |
| 4. | *β-actin*_RP | TGCCAACCATCACTCCCTGA1 | |
